# Caspase-8 deficiency induces a switch from TLR3-induced apoptosis to lysosomal cell death in neuroblastoma cell lines

**DOI:** 10.1101/2020.01.13.904987

**Authors:** Marie-Anaïs Locquet, Gabriel Ichim, Aurelie Dutour, Serge Lebeque, Marie Castets, Kathrin Weber

**Author notes:** shared last and corresponding authors.

## Abstract

TLR3 converts in cancer cells from an inflammatory to a death receptor and TLR3-induced cell death activates the extrinsic apoptosis pathway. Here, we demonstrate that activation of TLR3 triggers a form lysosomal cell death. Following the combinational treatment of IFN-1/poly(I:C) of the Caspase-8 deficient neuroblastoma cell line SH-SY5Y, lysosomes enlarge and accumulate before cells display characteristic apoptotic morphologies. However, caspases are not involved in signalling from TLR3 to the lysosome as 25 μM did not inhibit cell death. However, increasing zVAD concentrations to 50 μM which is known to inhibit cathepsins, as well as a specific cathepsin B inhibitor reduced TLR3-induced lysosomal cell death. Thus lysosomal cathepsins have a role in cell death execution and overtake the role of caspase-8 in inducing the apoptotic caspase cascade. Further in the caspase-8 positive neuroblastoma cell line SK-N-AS, knockdown of caspase-8 induces a switch from TLR3-induced apoptosis to lysosomal cell death. Taken together our data suggest that lysosomal cell death represents a default death mechanism, when caspase-8 is absent.

## INTRODUCTION

One of the hallmarks of cancer is the evasion of apoptosis ^1^ which results in resistance to targeted cell death based anti-cancer therapies. The therapeutic challenge lies within the bypass of the apoptosis-resistance of cancer cells by inducing alternative forms of programmed cell death. Next to apoptosis, two additional major forms of cell death exist, including autophagic/lysosomal cell death (Type II), and necrosis (Type III) ^2^. Both forms can be executed in a programmed manner, providing a possible cell death alternative to apoptosis in order to kill cancer cells. While necroptosis, a form of programmed type III cell death, is already discussed as an alternative for targeted cell death based therapies, the functioning of autophagy as an active cell death mechanism remains highly controversial. In general, autophagy is a survival mechanism to overcome cellular stress situations, like nutrient deprivation ^3^. Thus autophagy probably accompanies rather than promotes cell death in most scenarios and rarely represents a failed survival attempt ^4^.

Type II cell death is known to be associated with the appearance of the (autophago-)lysosomes, which do not only represent the degenerative endpoint of autophagy and endocytotic pathways, but lie at the crossroads of life and death pathways. As an acidic compartment with limiting membranes, lysosomes contain various types of proteolytic enzymes, like cathepsins, to facilitate recycling of cellular components and the breakdown of major macromolecules, including lipids, polysaccharides and proteins ^5^. During lysosomal cell death the lysosomal membrane permeabilises resulting in the release of lysosomal cathepsins into the cytosol. Once released lysosomal proteases can engage the mitochondrial pathway of apoptosis directly by processing of caspases or indirectly by for example the cleavage of Bid to truncated (t)Bid^6^. At the mitochondria tBID activates BAX/BAK leading to cytochrome c release and subsequent activation of the caspase-cascade with caspase-9 as the apical caspase. In addition also caspase-independent cell death mechanisms can be induced by LMP such as ROS an iron release ^7^. Although the exact underlying mechanism for LMP remains unknown, it can be induced by an increase in oxidative stress which might arise from the accumulation of lysosomes, as well as certain anti-cancer reagents^8^.

In contrast to apoptosis and necroptosis, lysosomal cell death is not directly triggered by the assembly of signalling platforms following activation of receptors of the Tumor Necrosis Factor Receptor (TNFR) family or Toll-like receptors (TLRs). TLR3 is an inflammatory receptor expressed on immune but also epithelial cells, with its synthetic ligands already used as adjuvant in cancer therapy due to this innate inflammatory signalling capacity^9^. Recently TLR3 was described to act as a death receptor exclusively in cancer cells, while sparing normal cells by inducing the extrinsic apoptotic pathway in an increasing range of different cancer types such as NB, hepatocharcinoma, lung cancer and mesothelioma ^10–13^. Thus TLR3 has the potential to act as novel therapeutic target to selectively kill a broad range of cancer cells. TLR3, in contrast to other members of the death receptor family signals from the lysosome, following its processing by cathepsins ^14,15^. The TLR3 death-signalling pathway induces the formation of an atypical signaling platform facilitating the activation of caspase-8 triggering the extrinsic apoptotic caspase-cascade ^16,17^. TLR3 might also induce the necroptotic pathway by recruiting and activating the necroptotic key factor RIPK3 to the TLR3 receptor complex. RIPK3 in turn phosphorylates the pseudokinase MLKL, which induces its translocation to the plasma membrane and causes its permeabilisation ^18–20^.

Until now lysosomal cell death and LMP is not associated with the activation of TLR3. Here we describe for the first time, that TLR3 induced cell death is executed by lysosomal cathepsins and is accompanied by the enlargement and accumulation of lysosomes, finally converging which converging in the activation of the mitochondrial apoptotic pathway. This cell death mode appears to be a default death mechanism, when Caspase-8 is not present and the extrinsic apoptotic pathway not be activated.

## RESULTS

### TLR3 induces receptor-mediated cell death in SH-SY5Y cells

TLR3 is known to convert from an inflammatory to a death receptor exclusively in cancer cells. In line Caspase-8 positive neuroblastoma cell line, SK-N-AS, was described that activation of TLR3 induces the induction of the extrinsic apoptotic pathway involving the activation of apical Caspase-8 ^10^. To investigate if TLR3 can induce an alternative programmed cell death pathway in the absence of Caspase-8, we chose the Caspase-8 negative neuroblastoma cell line SH-SY5Y. While SH-SY5Y cells are negative for TLR3 expression we pre-treated the cell line with IFN-1 for 16 h, which is known to upregulate TRL3 expression, followed by poly(I:C) treatment for indicated times **(Figure 1A)**. Following the combined treatment of IFN-TYPE I (IFN-1) and the synthetic TLR3 ligand poly(I:C), several TLR3 bands are apparent at 24 h and more pronounced at 48 h. Next to the full-length protein, represented by a 130kD band (glycosylated form) and a 100 kDa band (non-glycosylated from), the 70 kDa cleavage fragment is readily detectable **(Figure 1A)**. In correlation with the appearance of the 70kDa TLR3 fragment which represents the signalling-competent form of TLR3, IFN-1/poly(I:C) treatment induces time-dependent increase in cell death **(Figure 1B)**. Poly(I:C) can not only activate TLR3, but also the cytosolic RNA sensors MDA5 and RIG-I. In order to determine the specificity of the observed cell death to the activation of TLR3 we silenced the expression of TLR3 by siRNA. Following 72 h TLR3 expression was efficiently silenced determined by the absence of all three TLR3 forms on Western blot **(Figure 1C)**. In line at 72 h of TLR3 knock down the combined treatment of IFN-1/poly(I:C) resulted in reduced cell death induction **(Figure 1D)**. To further strengthen the dependency of the observed cell death on the activation of TLR3 poly(I:C) was replaced by poly(A:U), which has a higher specificity to TLR3 then to the cytosolic RNA sensors. In agreement with the TRL3 knockdown, treatment with IFN-1/poly(A:U induced cell death in SH-SY5Y cells at a comparable level than poly(I:C) **(Figure 1E)**. In order to provide evidence that observed IFN-1/poly(I:C) induced cell death is triggered locally at the activated TLR3 receptor and not through an TLR3-mediated autocrine TNF-α loop, the sensitivity of SH-SY5Y to TNF-α was determined. In agreement with the previous findings that SH-SY5Y cells do not express TNFR ^21^, no cell death was observed neither with or without IFN-1 pre-treatment **(Figure 1F)**. Taken together these results indicate that cell death induced by IFN-1/poly(I:C) treatment is specific to the activation of TLR3 and does neither involve the cytosolic RNA receptors, nor represents a secondary effect mediated by an autocrine TNFR activation.

**Figure 1.**
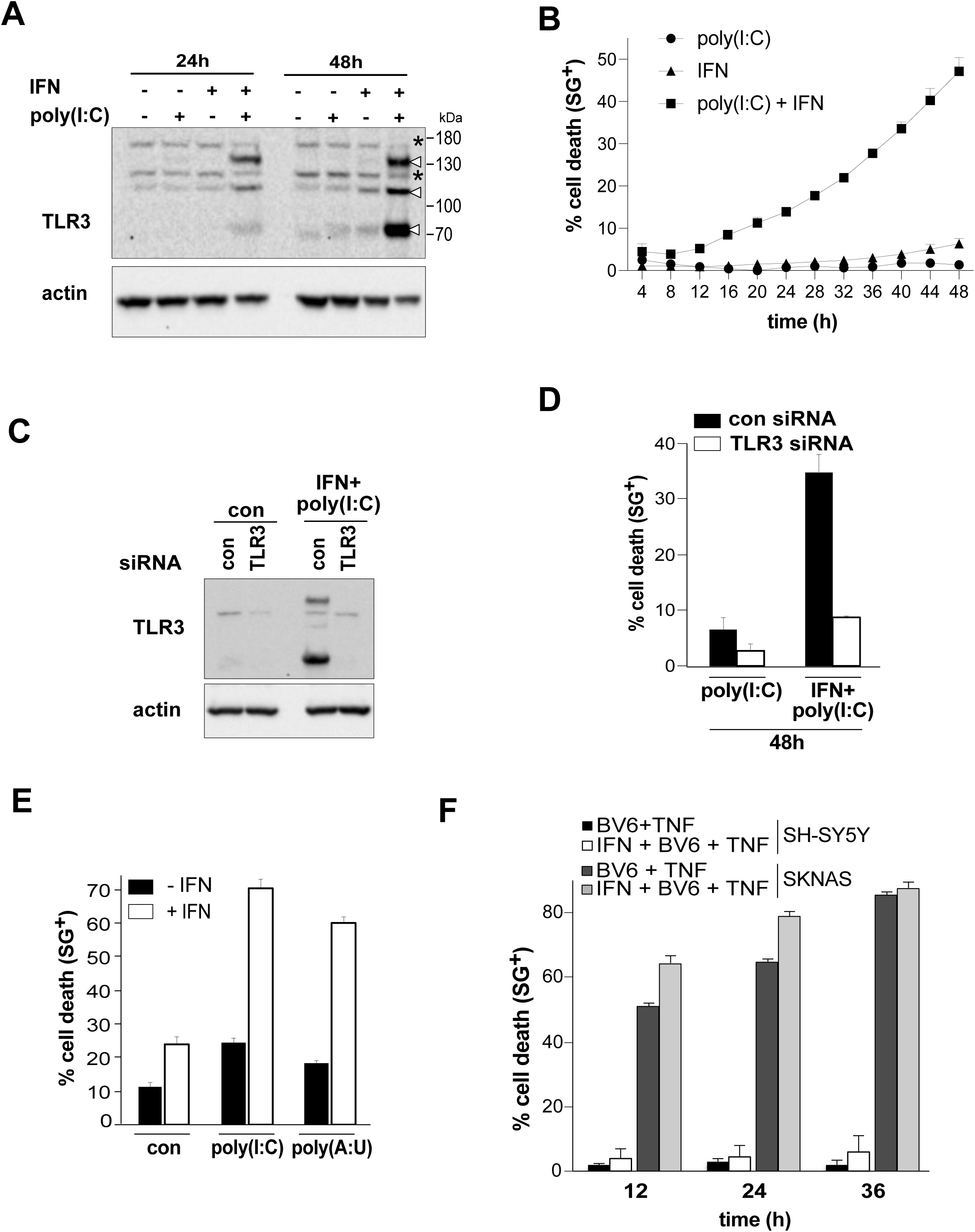
IFN-1/poly(I:C) treatment induces TLR3-specific cell death in caspase-8 negative SH-SY5Y. (A) TLR3 expression level of SH-SY5Y cells following IFN-1/poly(I:C) for 24 h and 48 h. (B) Cell death profile of SH-SY5Y pre-treated with IFN-1 for 16 h followed by poly(I:C) for the indicated time points. The percentage of SYTOX Green^+^ cells was analysed and profiles are averages ± S.E.M. *n* (number of independent experiments) = 3. (C) TLR3 expression levels of SH-SY5Y cells following siRNA mediated downregulation of TLR3 and treatment with IFN-1/poly(I:C). (D) Cell death profile of TLR3 siRNA transfected SH-SY5Y cells following IFN-1/poly(I:C) treatment for 48 h determined by Sytoxgreen uptake. Data points represent the average ± S.E.M of three independent experiment (n=3). (E) Cell death profile of SH-SY5Y cells treated with IFN-1/poly(I:C) or IFN-1/poly(A:U) for 48 h, determined by PI uptake. (F) Cell death profile of SH-SY5Y cells treated for the indicated times with BV6 + TNF or IFN-1 + BV6 + poly(I:C). Cell death was determined by Sytoxgreen positivity. Data points represent 3 independent repeats. SEM is indicated. SK-N-AS cells were used as a positive control.

### TLR3-induced cell in SH-SY5Y cells does not resemble classical apoptotic or necroptotic cell death

TLR3 activation induces in tumour cells the extrinsic apoptotic pathway, with caspase-8 being activated as apical caspase ^16,17^. While SH-SY5Y cells are categorized as caspase-8 negative cell line ^22^, it was described that IFN-γ can upregulate caspase-8 expression in SH-SY5Y cells, rendering cells sensitive to TRAIL-induced apoptosis ^23^. In analogy IFN-1 might upregulate caspase-8 expression sensitising SH-SY5Y cells to poly(I:C)-induced extrinsic apoptosis. However, IFN-1 did not upregulate Caspase-8 expression levels **(Figure 2A)**. Further, in correlation with no caspase-8 expression, IFN-1-treatment did not sensitised SH-SY5Y cells to TRIAL-induced apoptosis (**Figure 2B**). Thus IFN-1/poly(I:C) treatment did not appear to engage the classical extrinsic apoptotic signalling pathway, leaving a caspase-independent form of programmed cell death, namely necroptosis, as possible explanation. However, essential necroptotic key factor RIPK3 is not expressed in SH-SY5Y, neither with or without IFN-1 treatment (**Figure 2A**). In agreement with the absence of RIPK3, the necroptosis inhibitor GSK872’, which blocks RIPK3 activity, had no effect on poly(I:C)/IFN-1 induced cell death in SH-SY5Y cells, while markedly reducing necroptotic cell death in HT-29 cells **(Figure 2C)**. According to the expression profile of SH-SY5Y cells, which revealed not only the absence of RIPK3, but also caspase-8, the induction of caspase-dependent cell death seemed unlikely. However, we still could detect the processing of caspases following IFN-1/poly(I:C) treatment, including the activation of the initiator caspase-9 as well as the effector caspase-3 and the cleavage of its substrate PARP **(Figure 2D)**. Of note, the proforms of caspase-3 and PARP were still readily detectable, suggesting their incomplete processing. Extrinsic apoptosis signalling can be amplified over a mitochondrial signalling loop, while Caspase-8 mediated cleavage of Bid to truncated (t)Bid interconnects the receptor apoptotic signalling with the mitochondrial one. Strikingly, despite the absence of Caspase-8, IFN-1/poly(I:C) treatment decreased the full length form of Bid, indicating its cleavage to tBID **(Figure 2D)**. To further investigate the involvement of caspases in the IFN-1/poly(I:C) induced cell death, the pan-caspase inhibitor zVAD was included. At low concentrations of 25 μM, which are sufficient to inhibit the caspase activity, zVAD had now effect on cell death induction, leaving the exclusive role of caspases in the execution of this observed cell death unlikely **(Figure 2E)**. At higher concentrations zVAD inhibits un-specifically cysteine proteases, like e.g. the lysosomal cathepsins. Indeed at 50 μM as well as 100 μM zVAD reduced IFN-1/poly(I:C) induced cell death in a concentration-dependent manner.

**Figure 2.**
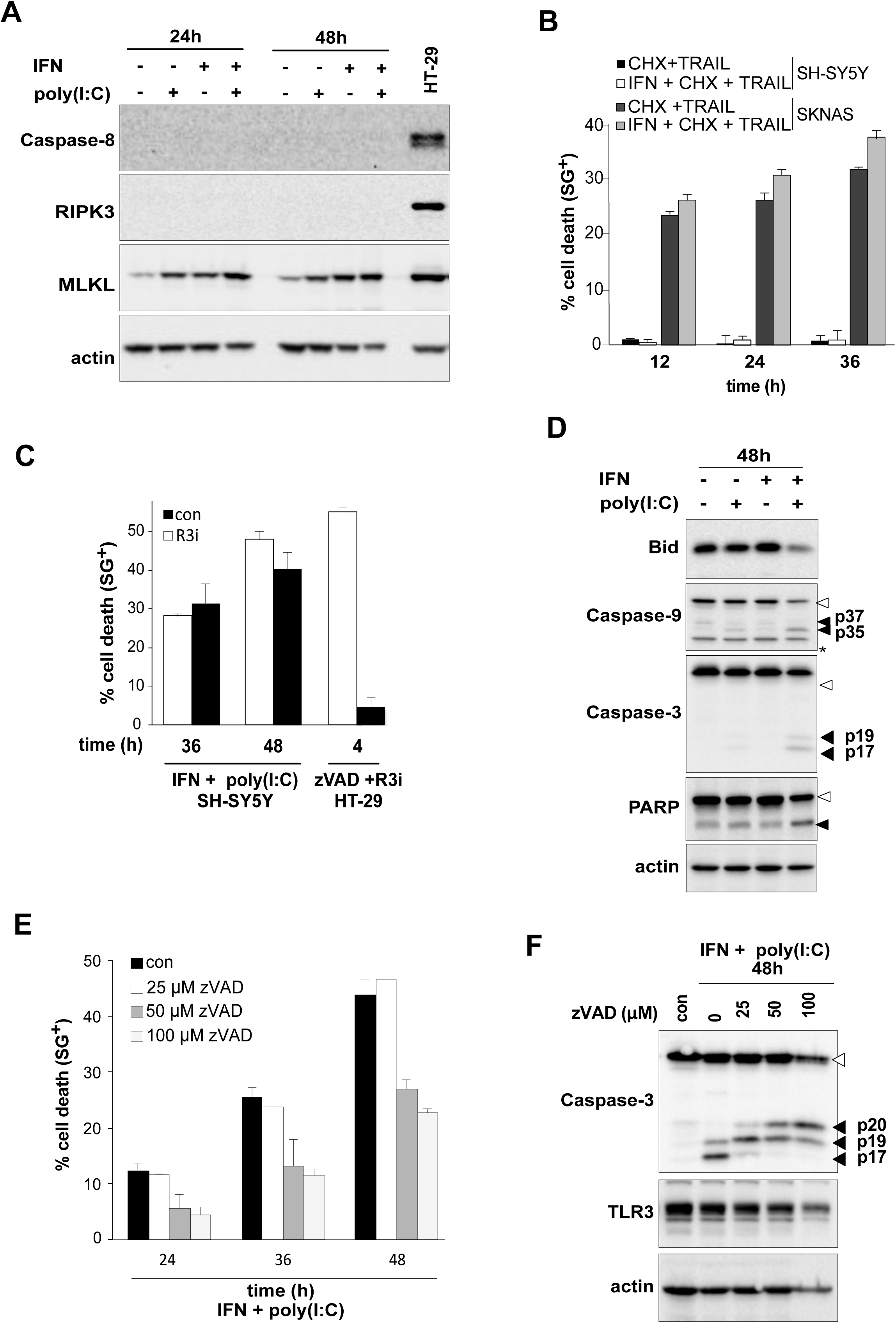
IFN-1/poly(I:C) induced cell death does not resemble extrinsic apoptosis nor necroptosis. (A) Expression levels of caspase-8, RIPK3 and MLKL in SH-SY5Y cells following treatment with IFN-1/poly(I:C) for 48 h. HT29 cells were used as a positive loading control. (B) Cell death profile of SH-SY5Y cells treated with IFN-1/TRAIL. The percentage of SYTOX Green^+^ cells was analysed and profiles are averages ± S.E.M. *n* = 2. (C) Cell death profile of SH-SY5Y cells pre-treated with the RIPK3 inhibitor GSK782’ followed by treatment with IFN-1/poly(I:C). Cell death was determined by Sytoxgreen uptake. Data points represent the averages ± S.E.M. of three independent experiments. HT-29 cells were used as positive control. (D) Western blot analyses of the processing of Bid, Caspase-3 and PARP following IFN-1/poly(I:C) treatment for 48 h. Proforms are indicated by an unfilled arrowhead, while the cleavage products are indicated by black arrowheads. Actin was used as a loading control. Astricks denotes unspecific band. (E) Cell death profile of SH-SY5Y cells pretreated with indicated concentrations of zVAD followed by IFN-1/poly(I:C) treatment. Cell death was determined by SytoxGreen positivity and datapoints represent the averages ± S.E.M. of n=3. (F) Caspase-3 cleavage as well as the expression levels of TLR3 of SH-SY5Y cells pre-treated with indicated zVAD concentration followed by treatment for 48 h with IFN-1/poly(I:C). Caspase-3 proform are indicated by an unfilled arrowhead, while the cleavage products are indicated by black arrowheads. Actin was used as a loading control.

To gain further insights into the effect of zVAD on IFN-1/poly(I:C) induced cell death, the processing of the effector caspase-3 was analysed **(Figure 2F)**. During its activation, caspase-3 is cleaved between the large and the small subunit, generating a p20 and p10 fragment. The p20 fragment is further processed by clipping off a pro-domain at the p20 fragment, resulting in the generation of a p19 and p17 fragment, respectively. Increasing concentrations of zVAD resulted in a shift from the p17 fragment to the p20 fragment, while at 100 μM zVAD the predominant fragment was the enzymatic less active fragment p20. Thus zVAD inhibited in a concentration-dependent manner the processing of caspase-3 while concentrations of at least 50 μM were required to inhibit, at least partially, the processing into p19 and p17 fragments. Importantly, TLR3 is described to be cleaved in the lysosomes by cathepsins to generate the signalling-competent variant of TLR3 which is characterized by a 70kDa fragment on Western blot ^24^. Increasing concentrations of zVAD, which are known to gradually inhibit cathepsins, did not prevent the generation of the 70 kDa TLR3 cleavage fragment **(Figure 2F)**. Thus decrease in cell death induction by increasing zVAD concentrations did not result from a block in the formation of signalling competent form of TLR3. Taken together these results suggest the involvement of cathepsins in the execution of IFN-1/poly(I:C)-induced cell death, while cathepsins could account for the cleavage of Bid to tBid which then activates the intrinsic apoptotic pathway with caspase-9 as the apical caspase.

### Impact of cathepsins and lysosomes in IFN-1/poly(I:C)-induced cell death

To provide further evidence for the involvement of cathepsins in IFN-1/poly(I:C) induced cell death the cathepsin inhibitor CATI-I, with the highest affinity against cathepsin B, the predominant cysteine protease in neurons. At 10 μM only a minor inhibition of IFN-1/poly(I:C)-induced cell death was observed (**Figure 3A**). In agreement with the general pro-survival role of cathepsin within the lysosomes, their inhibition by 25 μM of CATI-I resulted in the induction cell death. Despite exerting auto-toxic properties, 25 μM CATI-I reduced the IFN-1/poly(I:C)-induced cell death, further suggesting a role of cathepsins in the execution of cell death **(Figure 3A)**. Of note, the incomplete cell death inhibition by CATI-I points towards the involvement of other cathepsins or additional cathepsin-independent death mechanisms in IFN-1/poly(I:C)-induced cell death in SH-SY5Y cells. In correlation with the cell death profile caspase-3 processing shifted in a concentration-dependent manner from the p17 to the p19 fragment **(Figure 3B)**. Thus cathepsins are activating caspases during IFN-1/poly(I:C) induced death. Under steady-state conditions cathepsins are stored in lysosomes, but released during lysosomal cell death into the cytosol, which might be facilitated by an accumulation and enlargement of lysosomes. To investigate a lysosomal accumulation during IFN-1/poly(I:C)-induced cell death, cells were stained with LysoTracker and analysed by FACS. Indeed, an increase in LysoTracker staining was detected at 24 h of IFN-1/poly(I:C), while the cell population shifted about an entire log phase. Timelaps imaging revealed an increase not only in the quantity of lysosomes but also in the size of the lysosomes (movie in supplemental data). Cells with accumulated and enlarged lysosomes eventually rounded up and showed membrane blebbing, characteristic for apoptotic cell death (**Figure 3D**). Taken together, the activation of TLR3 induces in SH-SY5Y cells a particular form of cell death which is associated with the accumulation of lysosomes and their permeabilisation, which results in the release of cathepsins into the cytosol, where they activate caspases to induce apoptosis.

**Figure 3.**
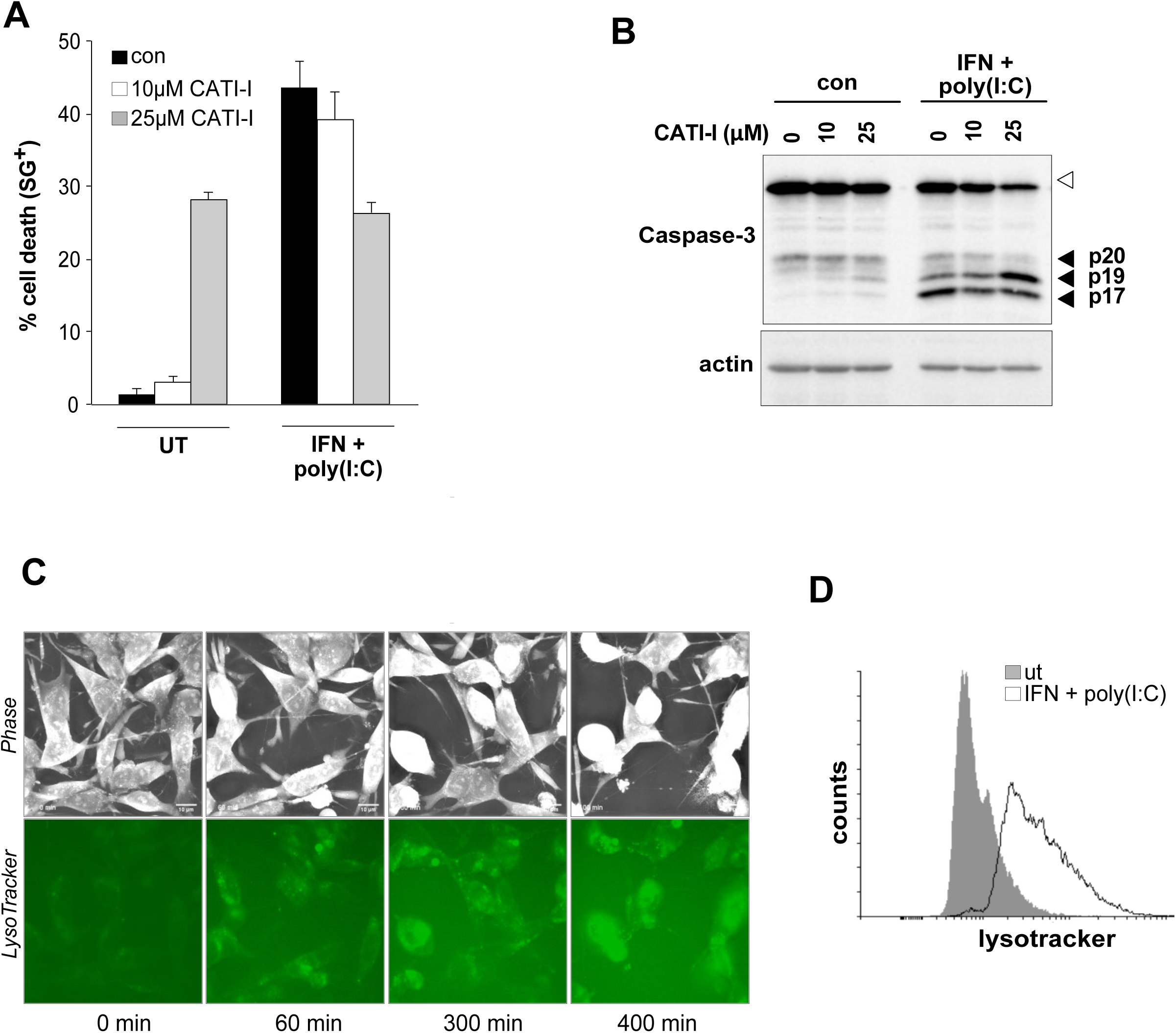
IFN-1/poly(I:C) induced cell death involves lysosomes. (A) Cell death profile of SH-SY5Y cells pre-treated with indicated concentration of the cathepsin inhibitor CATI-I followed by treatment with IFN-1/poly(I:C) for 48h. The percentage of SYTOX Green^+^ cells was analysed and profiles are averages ± S.E.M. *n* (number of independent experiments) = 3. (B) Cleavage of Caspase-3 of SH-SY5Y cells following treatment with indicated concentrations of CATI-I and IFN-1/poly(I:C) for 48 h. Caspase-3 proform are indicated by an unfilled arrowhead, while the cleavage products are indicated by black arrowheads. Actin was used as a loading control. (C) Time series of SH-SY5Y cells 2D Refractive index I maps, individual lysosome-specific fluorescence signal (LysotrackerGreen™), and overlay of Lysosome segmentations after treatment with IFN-1/poly(I:C) (OA) (related to movie shown in Supplemental information. (D) Histogram analysis of LysoTracker staining of con treated (gray filled histogram) overlayed with IFN-1/poly(I:C) (48 h) treated (black line) SH-SY5Y cells. Depicted histogram is a representative of n=3.

### Caspase-8 knock down induces a switch from TLR3-induced apoptosis to lysosomal cell death

Neuroblastoma, and in particular the therapy-resistant high-grade type, are characterised by a lack in caspase-8 expression resulting in apoptosis-resistance. To gain insights if the observed TLR3-induced lysosomal cell death is triggered in dependency to the presence or absence of caspase-8, we switched cell lines to the caspase-8 positive, TLR3 positive neuroblastoma cell line SK-N-AS. In correlation with earlier reports, SK-N-AS cells were sensitive to TLR3-induced apoptosis resulting in the activation of caspase-8 as an apical caspase^10^, **Figure 4A**. Of note, in our hands the combined treatment of the cIAP inhibitor BV6 and poly(I:C) is required to induce apoptosis in SK-N-AS cells, which probably results from culturing conditions between a minimal confluency of 40% and a maximal confluency of 70-80%. Knock down of caspase-8 by RNA interference resulted in sensitisation to TLR3 induced cell death **(Figure 4A)**. This result is in sharp contrast to our earlier finding that TLR3-induced cell death is nearly completely abolished by siRNA-mediated downregulation of caspase-8 expression in lung cancer cell lines ^17^. In addition, TLR3 activation resulted, in the activation of extrinsic caspase cascade in control conditions, with caspase-8 being the apical caspase followed by activation of caspase-3 which converges in the cleavage of PARP **(Figure 4B)**. However, when expression of caspase-8 was downregulated by siRNA, caspase-3 was still processed albeit less efficient then in control conditions, as the predominant fragment detectable was the catalytic inactive p20. Consequently also PARP processing was decreased compared to control as the proform was readily detectable. Thus these results suggest that knockdown of caspase-8 induces a switch from classical extrinsic apoptosis to a form of cell death in which the effector caspase-3 is activated by an alternative, caspase-8-independent mechanism. To investigate if cathepsins might activate caspase-3 we included the cathepsin inhibitor CATI-I into the cell death assay. Importantly, like in SH-SY5Y cells, treatment with CATI-I did not interfere with the generation of the 70 kDa signalling competent cleavage fragment of TLR3 **(Figure 4C)**. CATI-I did not alter cell death induction by IFN + BV6 + poly(I:C) in control conditions, but reduced cell death when caspase-8 was knocked down **(Figure 4D)**. Taken together these results point towards a switch mechanism in dependency to caspase-8, while in the presence of caspase-8 apoptosis, in the absence a cathepsin-dependent cell death occurs.

**Figure 4.**
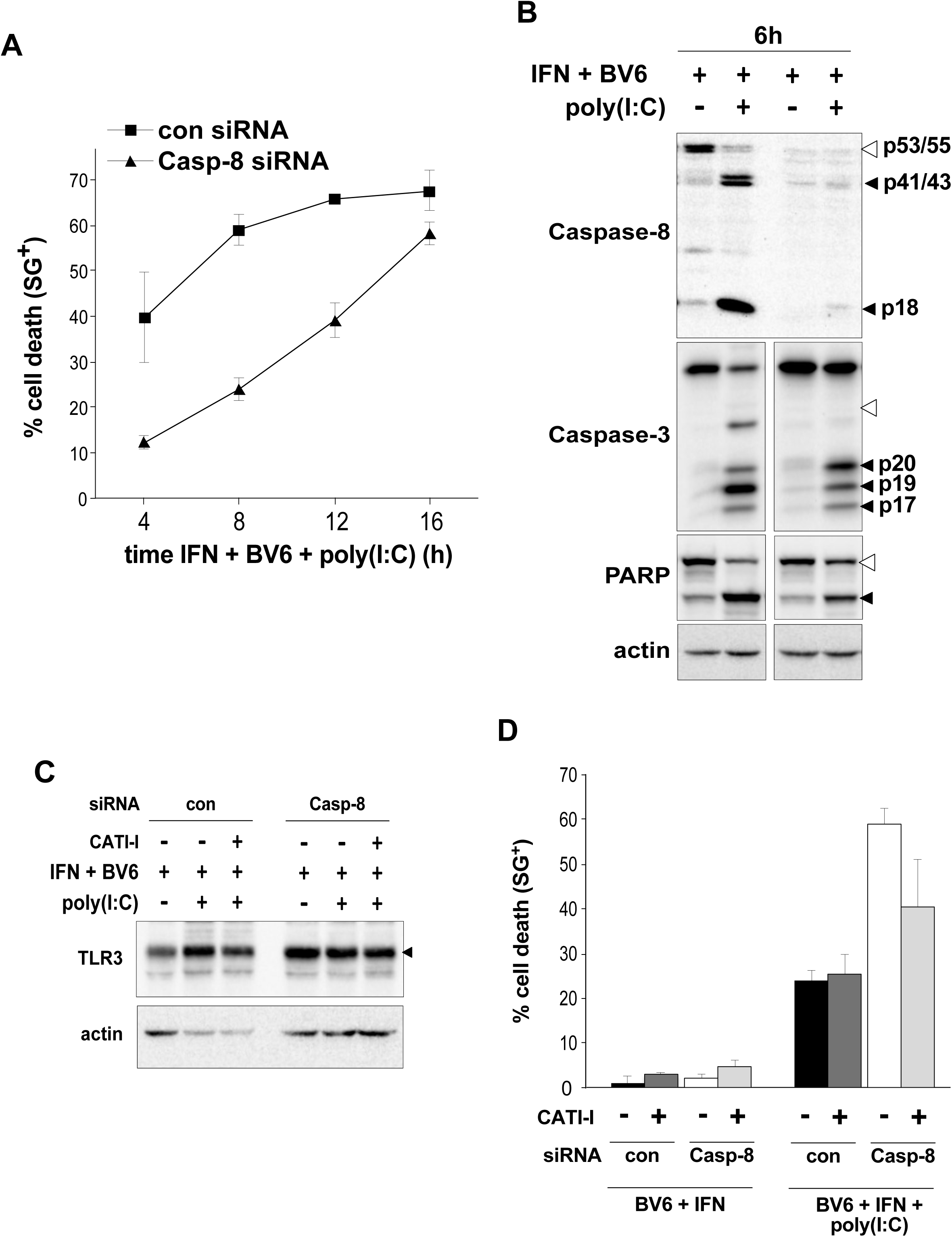
Knock down of caspase-8 induces a switch from TLR3-induced apoptosis to lysosomal cell death. (A) Cell death profile of SK-N-AS cells transfected either with control (con) or caspase-8 (Casp-8) siRNA for 72 h and treated with BV6 + IFN + poly(I:C). Cell death was determined by Sytox green uptake and data points represent the averages ± S.E.M. of at least three independent repeats. (B) Caspase-8, caspase-3 and PARP cleavage following BV6 + IFN + poly(I:C) treatment of SK-N-AS transfected with con or Casp-8 siRNA. Proforms are indicated by an unfilled arrowhead, while the cleavage products are indicated by black arrowheads. Actin served as loading control. (C) Expression levels of the signalling-competent cleavage fragment of TLR3 (depicted by a black arrowhead) of SK-N-AS cells transfected with con or Casp-8 siRNA and pre-treated with 10μM CATI-I followed by treatment with BV6 + IFN + poly(I:C). Actin was as a loading control. (D) Cell death profile of SK-N-AS cells transfected with con or Casp-8 siRNA and pre-treated with 10μM CATI-I followed by treatment with BV6 + IFN + poly(I:C). Cell death determined by Sytoxgreen positivity. Bars represent the averages ± S.E.M. of n=3.

## Discussion

Three different types of cell death exist, including apoptosis (Type I), autophagic cell death (Type II), and necrosis (Type III). While apoptosis is the main form of cell death which occurs during development and ensures tissue homeostasis, during the development of cancers different mechanisms are establish to evade apoptosis. For example, a common mechanism to achieve apoptosis resistance in neuroblastoma is the silencing the expression of caspase-8, the initiator caspase of the extrinsic apoptotic pathway. Further, the absence of caspase-8 does not only block the extrinsic apoptotic pathway, but also inhibits the activation of the intrinsic apoptotic pathway-resulting in the resistance to a wide spectrum of drugs and the failure of targeted cell-death based treatments. This is in particular true for stage 3/4, high grade neuroblastoma, which continue to have very poor prognosis and only 50 % of children with high-risk neuroblastoma can be currently cured. Moreover, intensive therapies administered during childhood are not devoid of long-term side effects, notably increasing life time risk for secondary malignancies. Hence, there is an urgent need for novel therapeutic strategies which bypass the apoptosis-resistance by inducing alternative programmed cell death pathways in order to combat neuroblastoma.

Our results reveal for the first time, that activation of the death receptor TLR3 can circumvent caspase-8 deficiency by inducing LMP, which leads to the release of lysosomal cathepsins into the cytosol. Although have optimal activity at the acidic pH of lysosomes, many lysosomal proteases can function at neutral pH of the cytosol ^8^. In this respect cathepsins were described to cleave Bid, providing a link between LMP and mitochondrial outer membrane permeabilisation^6^, Figure 2 or may cleave/activate caspases directly. Irrespective of the mechanism controlled LMP merges in most scenarios in the activation of caspases resulting in the induction of caspase-dependent cell death with apoptotic-like features like membrane blebbing (Figure 3). Thus cathepsins overtake the role of caspase-8 in activating the caspase cascade and can bypass the apoptosis-resistance of caspase-8 negative neuroblastoma cells.

LMP may also amplify or initiate cell death signalling in the context autophagy-dependent cell death^3^, we cannot answer the precise involvement of autophagy in this form of cell death yet. However, our results reveal that TLR3-induced cell death was accompanied by an increase of the number as well as the size of lysosomes (Figure 3). Such an increase in lysosomes could be explained by an increased autophagic flux, while autophagic lysosome reforming might be inhibited. Considering that both cell lines used in this study are neuroblastoma and as such derivate from the neuronal crest lineage attributing them with neuronal features. In particular neurons are characterized by high autophagic flux, which is further accelerated by the fact that both cell lines are cancer cells. Consequently these high levels of autophagy could promote the accumulation and enlargement of lysosomes resulting in priming lysosomes to LMP and sensitising the cells to TLR3 induced lysosomal cell death. Interestingly lysosomal membrane permeabilisation in the context of autophagy-dependent cell death was associated with the spontaneous regression of stage 4S neuroblastoma ^25^. Thus TLR3 induced lysosomal cell death in the context of autophagy might have the potential to induce tumor regression.

The induction of lysosomal cell death by death receptors was already described earlier, while for instance, TNF-α can induce LMP in a caspase-dependent manner through caspase-8 or in a caspase-independent manner involving RIPK1 and oxidative stress ^26^. In contrast our results reveal that activation of TLR3 induces caspase-independent LMP, while caspases are activated downstream of LMP explaining the observed apoptotic morphologic of the dying cell (Figure 3). Further this form of cell death seems to be default death mechanism for (neuronal) cells which lack the apoptosis machinery as evident by the switch from apoptosis to lysosomal cell death in response to caspase-8 knock down (Figure 4).

Taken together our results provide insights into an alternative programmed cell death pathway, which is triggered by the activation of TLR3. Importantly it does not require the presence of caspase-8 providing an alternative cell death pathway which can bypass the apoptosis-resistance of neuroblastoma. Although we are just starting to understand this cell death pathway TLR3-induced lysosomal cell death could therefore present novel therapeutic strategy to combat neuroblastoma.

## Supporting information

Supplemental Movie LysoTracker

## Methods

### Cell culture and treatment procedure

SH-SY5Y as well as SK-N-AS cells (both purchased from ATCC) were cultured in Dulbecco’s modified eagle’s minimal essential medium. HT-29 cells were purchased from ATCC and cultured in McCOY’s medium. Both media were supplemented with 10% fetal calf serum (FCS) and L-glutamine (200 mM). Cell cultures were routinely tested for mycoplasma contamination. SH-SY5Y cells as well as SK-N-AS cells were cultured between a minimal confluency of 40% and a maximal confluencey of 70-80%. To trigger TLR3-induced lysosomal cell death, SH-SY5Y and SK-N-AS cells were pre-treated for 16 h with 1000 Units/ml Interferon Type 1 (Interferon α/β, R&D Systems (Minneapolis, MN, USA)) followed by 2,5 μM BV6 (Selleckchem) pre-treatment (SK-N-AS cells only) and treatment with 10 μg/ml poly(I:C) (Invivogen). Human TNF-*α* (600 lU/ml) from R&D Systems (Minneapolis, MN, USA) was used at 50nM and human sTRAIL/Apo2L from PeproTech (Neuilly-Sur-Seine,France), which a pre-treatment with cyclohexamide (CHX, Sigma) for 2 h preceded. The following inhibitors of cell death were used; 25, 50 and 100 μM z-Val-Ala-DL-Asp(Ome)-fluoromethylketone (zVAD-fmk), 5μM RIPK3 inhibitor GSK872’ (both Selleckchem) and 10 and 25 μM Cathepsin inhibitor −1, CATI-I (Z-FG-NHO-Bz) (Calbiochem).

### Antibodies

The following antibodies (Ab) were used for Western blotting: anti-TLR3 (clones D10F10), anti-caspase-8 (clone 1C12), anti-human caspase-9, anti-caspase 3 (D3R6Y)), anti-PARP (46D11) and anti-Bid human-specific all from Cell Signaling Technology (Danvers, MA, USA). Anti-actin-HRP was purchased from Abcam.

### RNA interference

Control and human TLR-3 (L-007745-00-0005) and caspase-8 siRNA ON-TARGET SMARTpool were obtained from Dharmacon. Cells were transfected with the siRNAs using Lipofectamin RNAiMAX (Thermofisher) according to manufacturer’s instructions for 48 h (caspase-8) or 72 h (TLR3). The final siRNAs concentration was 20 nM for Caspase-8 and 40 nM for TLR3.

### Cell death assay

Cells were stained with 5 μM SYTOX Green (Life Technologies) and analysed by using a FLUOstar Omega fluorescence plate reader (BMG Labtech GmbH, Ortenberg, Germany). Percent cell death was calculated as follows:

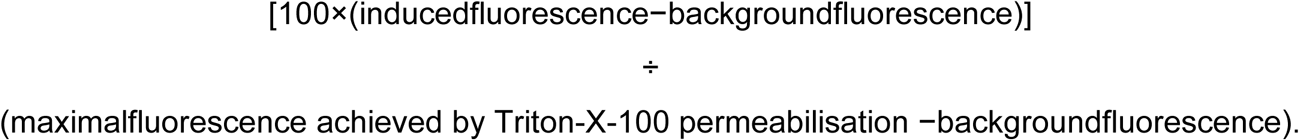

The data are presented as mean ± S.E.M of at least 2–3 independent experiments. All cell death profiles in this paper are shown as averages ± S.E.M.

### Holo-tomographic microscopy (HTM)

SH-SY5Y cells were seeded on Fluorodishes (ibidi GmbH, Gräfeling, Germany) and stained with 50nM LysotrackerGreen™ (Thermo fisher) 15 min prior imaging according to manufacturer’s protocol. HTM, in combination with epifluorescence, was performed on the 3D Cell-Explorer Fluo (Nanolive, Ecublens, Switzerland) using a 60× air objective at a wavelength of λ = 520 nm. Physiological conditions for live cell imaging were reached with a top-stage incubator (Oko-lab, Pozzuoli, Italy). A constant temperature of 37°C and an air humidity saturation as well as a level of 5% CO_2_ were achieved throughout the acquisitions. Refractory index maps were generated and every 5 min. Images were processed with the sofware STEVE.

